# Achieving natural behavior in a robot using neurally inspired hierarchical control

**DOI:** 10.1101/2021.01.22.427862

**Authors:** Joseph W. Barter, Henry H. Yin

## Abstract

Terrestrial locomotion presents tremendous computational challenges on account of the enormous degrees of freedom in legged animals, and the complex and unpredictable properties of the natural environment and the effectors. Yet the nervous system can achieve locomotion with ease. Here we introduce a quadrupedal robot capable of goal-directed posture control and locomotion over rough terrain. The underlying control architecture is a hierarchical network of simple negative feedback control systems inspired by the organization of the vertebrate nervous system. Without using an internal model or feedforward planning, and without any training, our robot shows robust posture control and locomotor behavior in novel environments with unpredictable disturbances.

## Introduction

The generation of adaptive movements is arguably the most fundamental function of the nervous system (17), and the underlying mechanisms remain poorly understood, especially for legged vertebrates with complex multi-jointed bodies (16). Movement cannot simply be equated with muscle output. Rather muscle outputs must vary by precisely the correct amount to counter the effects of continuously varying and unpredictable disturbances (10, 15, 42, 47, 51). To generate behavior through forward computation represents a very difficult computational problem for the nervous system: it requires accurate prediction of the required neural output to successfully produce any behavior while accounting for imperfect sensors, changes in muscle properties over time, a high level of redundancy in effectors, and unpredictable environmental conditions. This problem is made exponentially harder as the degrees of freedom are increased, a problem known as the “curse of dimensionality” (48).

There is an alternative approach that does not require forward computation in the first place. According to this perspective (39, 40), motor behavior may be viewed as the process of continuously controlling *perceptual inputs* through negative feedback. Because the system is controlling its perceptual input rather than its behavioral output, sensory perceptions are simply compared to changing internal goals and the errors between them drive corrective outputs that influence those perceptions. For such control systems to deal with the challenges of the natural environment, however, a hierarchical organization is required. In such a hierarchy, higher level controllers achieve their own reference states by adjusting the reference states of lower systems.

Perceptual variables such as muscle tension and length are thought to be controlled at lower levels in the spinal cord, while progressively more abstract variables including body posture, orientation, and position in space are controlled at higher levels in the brainstem, midbrain and basal ganglia (51). By utilizing this hierarchical network approach, highly complex goal-directed behavior can be decomposed into a set of simple nested control functions operating in parallel.

To test whether such a hierarchical organization can successfully generate natural behavior, we developed a quadrupedal robot with a hierarchical perceptual control architecture. This robot is capable of generating posture control, locomotion, orientation and basic navigation over uneven terrain. The ‘nervous system’ of the robot operates through a hierarchical network of simple control system modules. Unlike other robot control architectures that perform model-based control and planning (11, 12, 13, 14, 22, 23, 25, 33, 45, 50), our control architecture generates robust and adaptive goal-directed behavior through a simple feedback process requiring no model of the environment, prediction of future states, or learning. Unlike architectures in which behavior is generated by environmental stimuli or internal system dynamics (1, 2, 3, 4, 18, 24, 28, 29, 32, 36, 35, 37, 43, 46), our architecture generates adaptive behavior by automatically achieving a set of continuously changing internal goals in a hierarchy. The only computations required are operations like subtraction and integration, which can be successfully implemented by analog computing devices.

## Results

The robot is a quadruped with 12 degrees of freedom in four three-jointed legs (**Fig 1A**). Each joint is equipped with an analog torque sensor and an analog joint angle sensor. Each foot has a ground reaction force sensor, and the trunk houses an inertial measurement unit (IMU) for sensing tilts of the body (**Fig 1B**).

**Figure 1.**
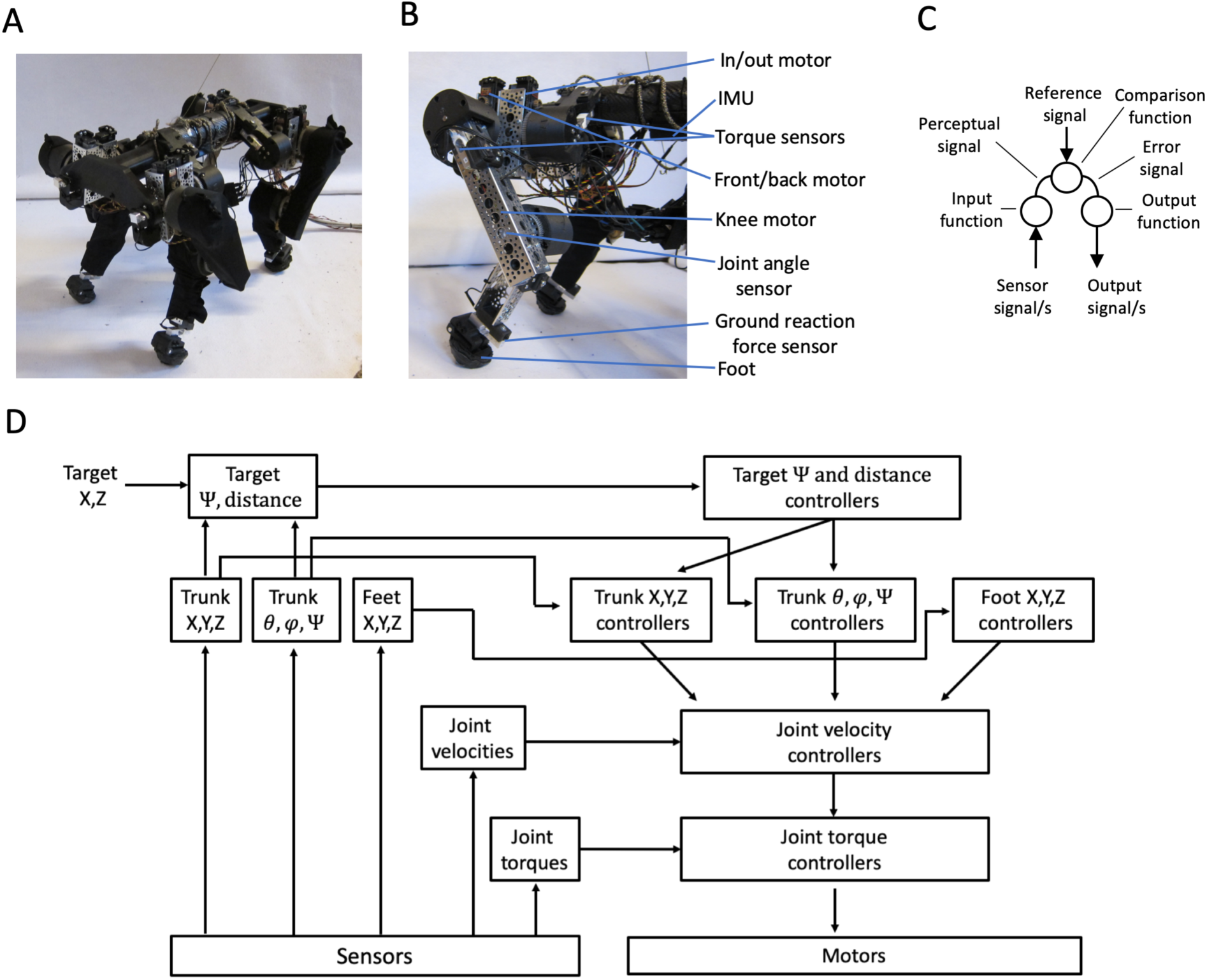
System overview. **(A)** Image of robot standing with neutral posture **(B)** Skin cutaway of a single leg showing leg parts. For each leg there are three joint angle sensors, and three torque sensors; one or each joint. Also shown is the inertial measurement unit (IMU) for the body. **(C)** Top: A simplified schematic of the closed loop control system module: a signal or signals are read into the input function where they are converted into a single perceptual value, which is compared with an internal reference value to generate an error. The error is converted by the output function into an output signal, which is sent out to a lower system. **(D)** Block diagram showing signal flow in our control architecture. Sensor inputs are converted by perceptual functions into joint torque perceptions, joint angular velocity perceptions, trunk X,Y,Z position and pitch(*θ*), roll(*φ*) and yaw(*Ψ*) perceptions. Yaw orientation to target is determined from target X,Z position and the trunk yaw perception. Distance to target is determined from target X,Z position and trunk X,Z position. Target position here is set by the experimenter.

All control systems at every level of the hierarchy have the same basic design. They vary only in their output tuning, output conversion functions and routing, and in the nature of the perceptual signal that each system controls. Each controller has three basic parts (**Fig 1C**): 1) an input function which converts raw sensory input values into a single perceptual value, 2) a comparison function which compares this perceptual value with a reference value coming from higher in the hierarchy for that perception and calculates the error between them, and 3) an output function which applies proportional-derivative (PD) tuning and a nonlinear conversion function (when necessary) to the error, and routes the resultant output signals to become reference signals for the appropriate set of lower level control systems. Control system modules in our robot were tuned manually by adjusting the PD gains of their output functions. Note that in all controllers, the variable being controlled is always the perceptual variable from the input function and *not* the output. The output of the system will vary according to both internal goals and perceptual inputs.

Sensory inputs are read, and outputs are commanded, in a continuous loop operating at a frequency of ∼230 Hz. During each iteration of the loop and starting at the top of the hierarchy, the perceptual value for each control system module is compared with a corresponding reference value to compute an error signal. The error for each active control system is then converted into the appropriate set of reference values for controllers one level down in the hierarchy. At the lowest level, errors are converted into output commands for the motors (**Fig 1D**). In this robot, behavior of the highest level controllers at the fourth level is determined by a sequence of locomotor target positions specified by the experimenter.

### Network levels 1-2: joint controllers

The lowest level in this control hierarchy is the level of joint torque control. For each joint in the body, torque is sensed and controlled by a dedicated controller. As with muscle tension control in a real animal, torque controllers may have a ‘ceiling’; torque inputs beyond these safe limits will cause the joint to yield, protecting it from damage. For the purposes of locomotor behavior, joint torque perceptions are individually normalized so that a value of zero corresponds to the baseline value measured while standing.

At the second level of the hierarchy, joint angular velocity is controlled. For each joint, angular velocity is sensed and compared to the reference angular velocity. The angular velocity error is converted into a reference signal for the corresponding joint torque controller at level 1. The angular velocity error and torque reference are both reduced through movement commanded by the torque controller,.

This velocity-torque hierarchy enables a form of viscous damped compliance; with a fixed angular velocity reference signal, any outside forces acting upon the joint will cause it to yield with a resistance roughly proportional to the velocity of the disturbance, as the angular velocity error is converted into a torque reference signal to oppose that error. Thus, the angular velocity controller gain determines the level of damping. This feature enables movements with the stabilizing and protective benefits of damped compliance. This feature was partly inspired by the lowest circuits in the spinal cord, which are responsible for controlling muscle tension and velocity. These circuits, which include type 1a fibers from muscle spindles and type 1b fibers from Golgi tendon organs, form a cascade control hierarchy (39).

### Network levels 3-4: posture and locomotor controllers

At the third level in the hierarchy, Cartesian positions and orientations of body parts are perceived and controlled. Because the lengths of body segments are known, we can convert a vector of joint angle measurements into perceptions of body position in 3D space through simple trigonometry. With an inertial measurement unit (IMU) to sense the trunk tilt angle, it is possible through a reference frame rotation to perceive body position in space relative to gravity. Foot ground reaction force sensors are used to detect ground contact.

As in the lower controllers, controllers at this level operate by computing the difference between reference and perception. Errors are then converted into the appropriate set of reference signals for joint angular velocity controllers that reduce those errors over time. The output function of a posture controller converts a single error into a set of reference signals for a group of lower controllers (**Fig 2A-B**).

**Figure 2.**
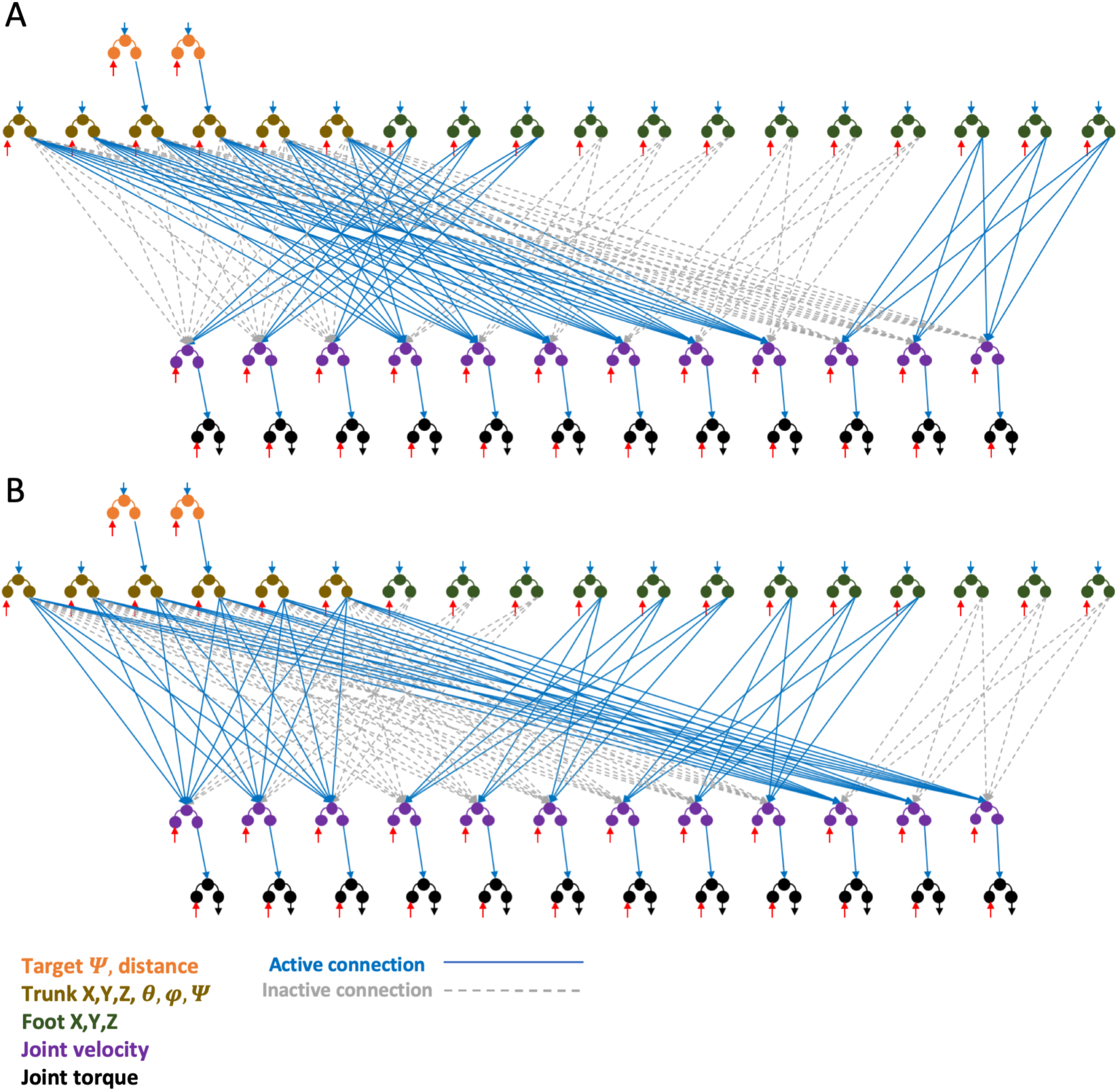
Network architecture diagram illustrating antagonism during stepping. Top and bottom diagrams both show the same network architecture. Blue lines represent active reference signals, red lines represent perceptual inputs and dashed gray lines represent inactive reference signals. The top **(A)** diagram shows active and inactive network connections during locomotion when the FL and BR feet are up and FR,BL feet are down. The joint velocity reference signals of the down legs are determined by the trunk orientation and position controllers, while the joint velocity reference signals of the up legs are determined by the foot position controllers. Orange controllers at level 4 represent target yaw (*Ψ*) orientation and position controllers. Brown controllers at level 3 represent trunk pitch (*θ*), roll (*φ*), and yaw (*Ψ*) orientation and X,Y,Z position controllers. Green controllers at level 3 represent foot X,Y,Z position controllers. Purple controllers at level 2 represent joint angle velocity controllers. Black controllers at level 1 represent joint torque controllers. The bottom **(B)** diagram shows the same thing as in (A) except with the leg pairs reversed.

Because the conversion between a given movement in Cartesian space and the corresponding joint angles is non-linear and depends on the current body posture, movements can be stabilized by including a simple gain scheduler in the output function which takes the current body posture into account. This gain scheduler preserves a linear relationship between error and output regardless of the current body posture. Otherwise, the output of a Cartesian or orientation controller is converted into a set of reference signals for joint velocity controllers according to fixed gains, resulting in movements that would be too large in certain postures and too small in others. For example, trunk height control in crouched vs. tiptoe postures require different joint angle changes to accomplish the same amount of vertical movement. The schedulers work by dynamically adjusting the effective gain of each output signal from a given level 3 controller to lower joint angular velocity systems so that the output is appropriately scaled for the current instantaneous posture and velocity of movement. The gain scheduler is a part of the output function of each level 3 postural controller.

Two types of perceptual variables are controlled at level 3: the positioning of each foot, and the positioning of the trunk. These controllers are antagonistic: the trunk controllers only operate through a given leg when the foot is planted on the ground, while the foot controllers only operate through that leg if the foot is unweighted and free to move about (**Fig. 2A-B**). For each leg there is one foot position controller for each of the three Cartesian axes of movement in the world reference frame relative to the vector of gravity. The Y axis controller perceives the position of the foot relative to the previous ground surface, while the X and Z axis controllers perceive the position of the foot relative to its position during neutral standing posture. Each foot position controller works by converting its error into three reference signals: one for each joint velocity controller of the same leg (**Fig. 3A**). When X, Y and Z controllers for a foot are running with fixed reference signals, then the foot remains fixed in space despite unpredictable trunk movements (**Fig. 3B**). Voluntary movement of the foot through changing position reference signals is also stable and exhibits good control (**Fig. 3C**), even when the body is also being tipped unpredictably (**Fig. 3D**). This capacity is important during locomotion for preventing feet from jamming into the ground when the trunk is disturbed.

**Figure 3.**
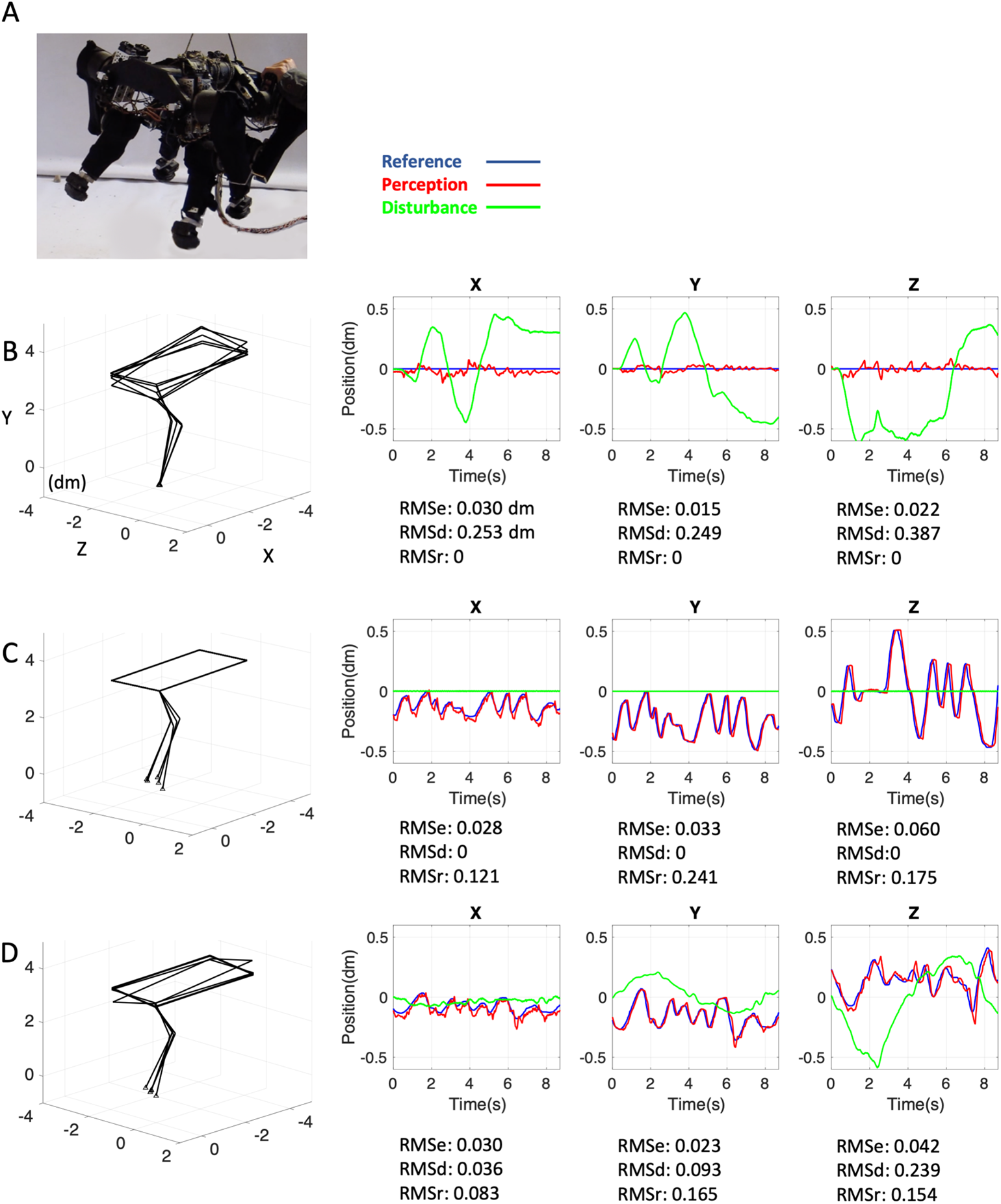
Foot position control. **(A)** The robot is shown hanging so that the trunk can be tipped while the legs are free to move. **(B)** Representative plots of X, Y and Z position control in the front left (FL) foot of a hanging robot with fixed X,Y and Z position reference signals while the experimenter randomly tips the body. Red lines represent perceptual signal, blue lines represent reference signal, and Green lines represent the disturbance, measured as the position of the foot due to body tilt if the controllers were off and the leg were rigid. For each plot, root mean squared error (RMSe), root mean squared disturbance (RMSd), and root mean squared reference signal (RMSr) values are reported, showing that the error is held small despite large changes in the disturbance. Wireframe snapshots illustrate foot stability, matching the stable reference signal, despite disturbances causing body movement. **(C)** Same plot format as in (B), showing a scenario in which the robot is not tipped but the foot position reference signals are randomly changed by the experimenter. RMSe is held at a low value despite large changes in RMSr. **(D)** Same plot format as above, showing a scenario in which the trunk is randomly tipped while the foot position reference signals are also randomly changed. RMSe is held at a low value despite large changes in both RMSd and RMSr.

For trunk orientation, there are 3 controllers: pitch, roll and yaw. Pitch and roll are perceived directly from the IMU, while yaw is perceived through integration of trunk yaw velocity relative to the feet that are planted on the ground. Each trunk orientation controller operates through legs that are planted on the ground; if 4 legs are planted, then the controller will adjust reference signals of 12 lower joint velocity controllers, whereas if 2 legs are planted, then the controller will adjust only 6 controllers (**Fig 4A**). Trunk orientation control can maintain its reference state when the ground is being unpredictably tipped (**Fig 4B, Movie 1**), when the ground is unmoving and the trunk is being voluntarily tipped through reference signal change (**Fig 4C, Movie 2**), or when the ground is being tipped while the trunk is being voluntarily tipped at the same time (**Fig 4D**). Pitch and roll reference signals were left at neutral values. During traverses of steep terrain with a slope greater than ∼15 degrees, trunk pitch is perceived relative to the pitch of the ground surface as sensed by the feet so that the trunk remains parallel with the ground.

**Figure 4.**
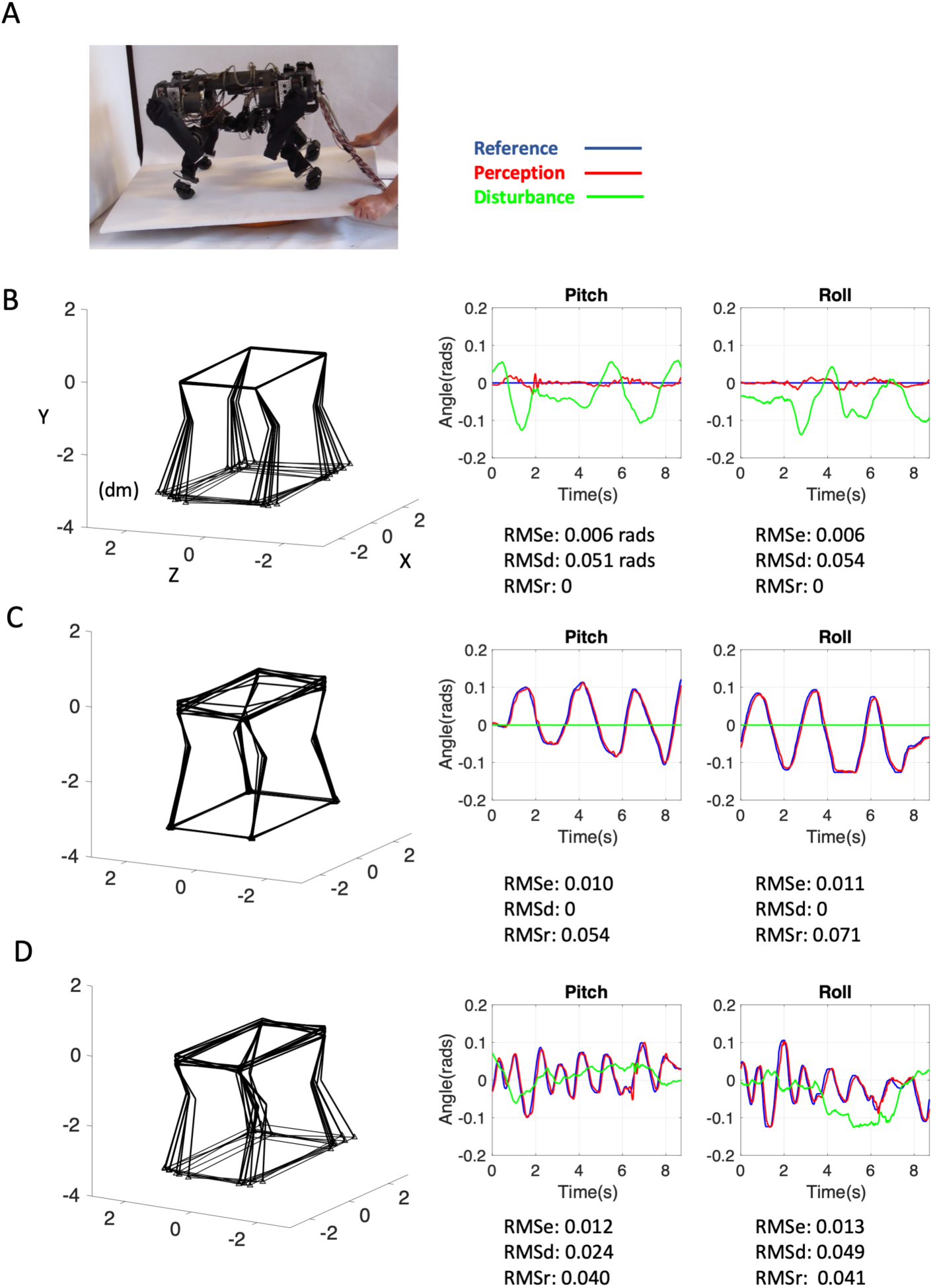
Trunk orientation control. **(A)** The robot is shown standing on a tippable platform. **(B)** Representative plots of trunk pitch and roll control while the experimenter randomly tips the platform. Red lines represent perceptual signal, blue lines represent reference signal, and Green lines represent the disturbance, measured as the angle of the platform. For each plot, RMSe, RMSd and RMSr values are reported, showing that the error is held small despite large changes in the disturbance. **(C)** Same plot format as in (B), showing a scenario in which the robot is not tipped but the pitch and roll reference signals are randomly changed by the experimenter. RMSe is held at a low value despite large changes in RMSr. **(D)** Same plot format as above, showing a scenario in which the platform is randomly tipped while the pitch and roll reference signals are also randomly changed. RMSe is held at a low value despite large changes in both RMSd and RMSr.

For trunk position control at level 3, there are three controllers: X, Y and Z. Trunk Y height is perceived by comparing the trunk position along that axis to the average position of the feet that are planted on the ground. X and Z are perceived as the path-integrated position of the trunk relative to the legs that are and have been planted on the ground. As with the trunk orientation controllers, these trunk position controllers operate through whichever legs are on the ground, so that the trunk can be maneuvered while the feet do not move. Combined, each down leg expresses the added output of all 6 trunk controllers. **Movie 3** shows control of trunk height and yaw while the reference signals are arbitrarily changed by the experimenter.

Rhythmic stepping up and down is accomplished by a simple burst generator for each leg. There is one burst generator per leg. When activated, a burst generator will switch the leg from trunk control to foot position control at the same time as it adjusts the Y foot position reference signal along a fixed path so that the foot lifts up and then back down and into the ground. Upon contact with the ground after achieving lift height, the burst generator resets and the leg switches back to effecting trunk control (**Fig 5A**). Because of motor speed limitations, achieved step height during locomotion was typically below 2 cm, but this value could be increased significantly with improved motors. Burst generators are activated in diagonal pairs so that the termination of both burst generators of a diagonal pair activates those of the opposite pair. Using this scheme, simply changing yaw orientation (**Fig 5B, Movie 4**) and trunk X position (**Movie 5**), reference signals during stepping result in locomotion. Trunk Y and Z position reference signals stay at a neutral zero value. The robot walks slowly, at approximately 8 cm/s. It is important to note that the low speed is due to the limited motors and 3D printed components, and does not reflect limitations in the design of the system architecture. When in swing mode, a foot will reach for a new X, Z reference position reflecting the velocity of the trunk, so that the feet step to continuously lead trunk movement, whether during voluntary locomotion or in response to an environmental disturbance (**Movie 6**).

**Figure 5.**
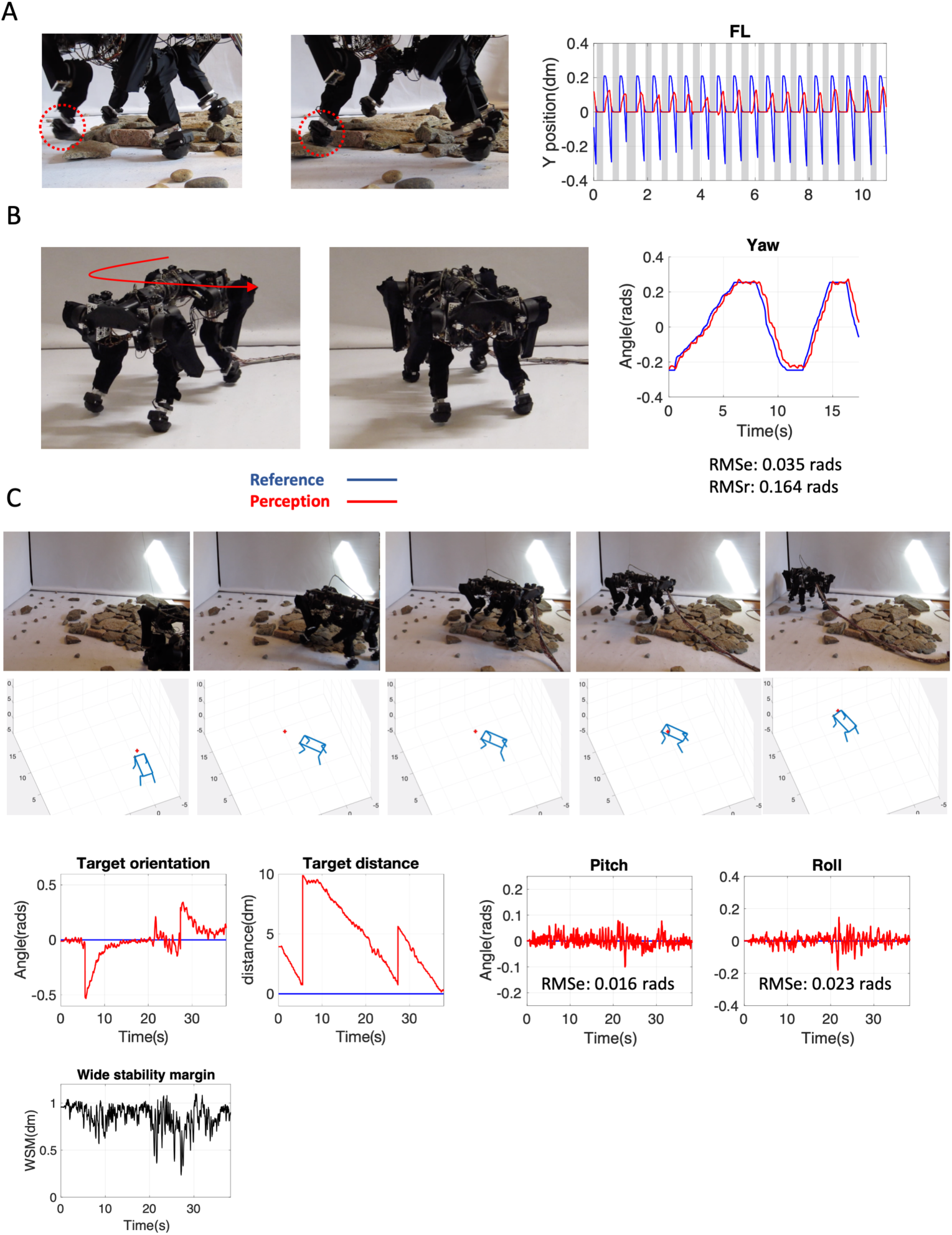
Stepping, orientation, and locomotion over rough terrain. **(A)** During stepping, the feet lift in alternating diagonal pairs. Two images show snapshot of the two major phases of the gait. The left image shows the stage in which the FL and BR legs are up and FR and BL legs are down. The right image shows the opposite stage. Plot shows representative plots of Y (distance from ground) position reference signals (blue) and perceptual signals (red) during stepping for the front left foot during stepping. Grey bars indicate when a given foot controller is off, and the leg is being used as part of the trunk control output functions. **(B)** During stepping, adjusting the trunk yaw reference signal causes the body orientation to change as the feet lift and reset position. **(C)** During stepping, the robot can successfully traverse rough and unstable terrain in pursuit of targets. A target orientation controller at level 4 adjusts the yaw reference signal at level 3, while a target distance controller at level 4 adjusts the front-back trunk position reference signal at level 3. Images are snapshots from a single target pursuit session. The location of the target at the time of each still image is illustrated in the corresponding wire frame diagram below as a red dot. Plots show target orientation, target distance, trunk pitch, trunk roll, and wide stability margin (WSM) measurements for the same session. Because of speed limits on locomotor behavior, orientation and distance errors in these systems reduce relatively slowly compared to lower systems, as illustrated in the target orientation and target distance control plots. The moment of sequence turnover can be seen in the plots as sharp increases in error. Pitch and roll plots illustrate that trunk orientation is continuously defended. WSM is calculated as the distance from the center of gravity to the nearest edge of the support polygon (Kimura, 2007). WSM plots illustrate that the center of gravity is successfully maintained above the support polygon.

At the fourth level in the hierarchy, orientation and distance to any specific location in the environment are controlled through locomotion. Since the robot is blind and has no distal sensors, position and orientation in the world are perceived by continuously integrating yaw velocity and translation velocity of the trunk by the down legs. Orientation relative to an external target is controlled by converting error into trunk yaw reference signal at level 3, while distance to the target is controlled by converting error into X (front-back) trunk position reference. In level 4 controllers of this robot, the reference signals are left at a neutral zero value so that the robot always orients towards a target and seeks to reduce target distance to zero. In these controllers, the target location is defined as part of the perceptual function and targets locations are sequenced by the experimenter.

Level 4 outputs are integrated so that they are converted into gradual changes in level 3 reference signals, and thus gradual changes in actual position and orientation. In this way, robot speed and orientation velocity are proportional to level 4 outputs. As shown in **Fig 5C**, by controlling trunk position and orientation relative to targets in the world, the robot is able navigate a path through the environment according to an experimenter-defined sequence of target position references. The robot is able to successfully reach each target in a 3-step sequence over flat terrain (**Movie 7**), and over uneven and unstable terrains including loose rocks (**Movie 8**) and rock piles (**Movie 9, Fig 5C**). **Fig 6A** shows several different terrains and corresponding target approach signals. During this behavior, the trunk controllers not only generate locomotor movement itself, but also defend against disturbances caused by external sources such as an experimenter push **(Fig 6B)**, as well as by terrain and the continuous disturbances caused by stepping itself **(Fig 6C)**.

**Figure 6.**
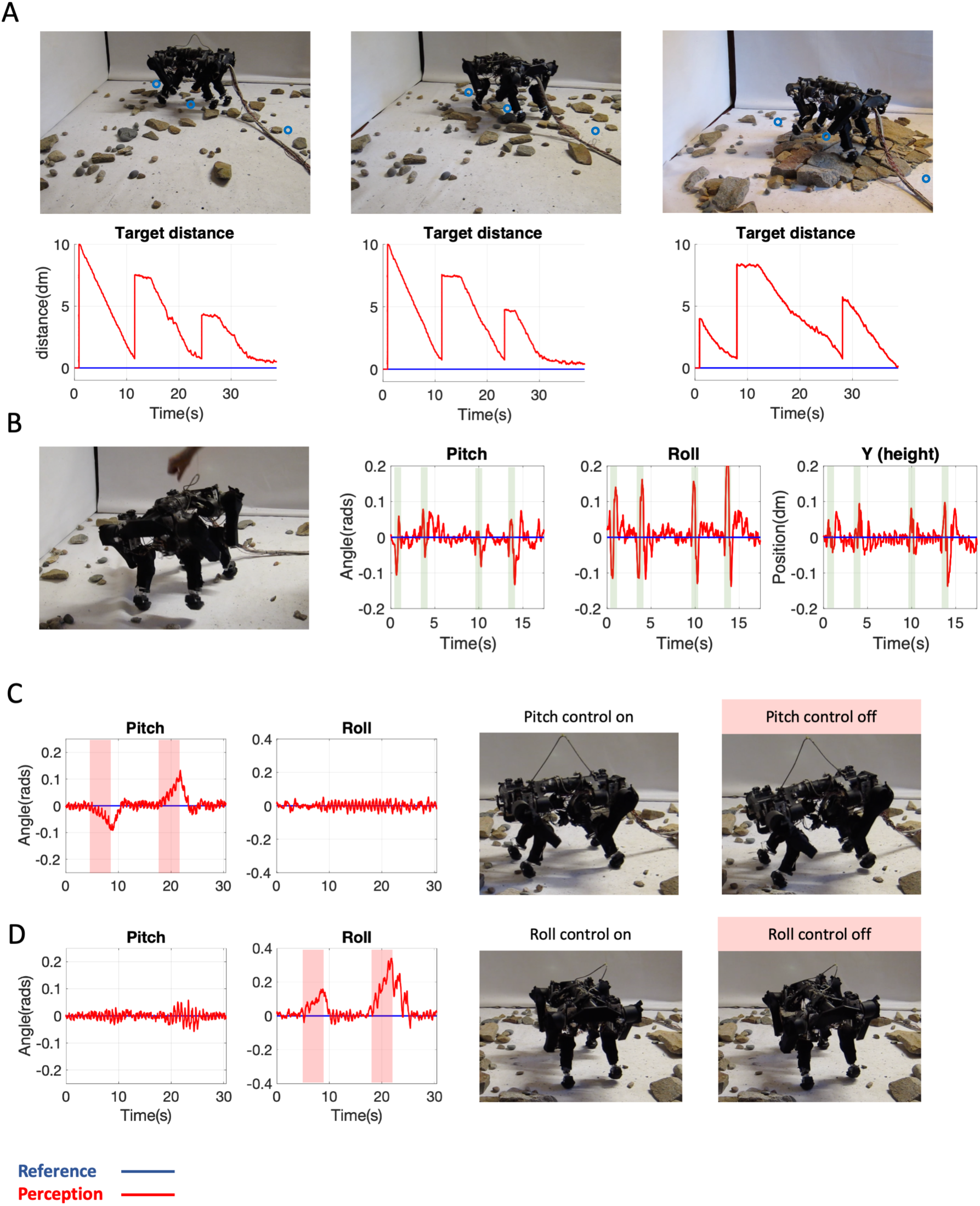
Locomotor control. **(A)** Three snapshots and corresponding target distance plots for locomotion to three targets across different rough terrains. Blue circles in the snapshots indicate the approximate location of targets. As shown in the plots, the robot hits all three targets by reducing the target distance error towards zero. **(B)** A snapshot shows the experimenter pushing the robot on its side, while plots show pitch, roll and trunk height control signals during repeated push disturbances. Red lines are perceptual signals, blue lines are reference signals, and push disturbances are indicated by light green bars. The plots show successful defense of the reference orientations and heights. **(C)** Plots showing perceptual and reference signals during a session in which the robot was walking in place when the pitch controller was periodically turned off, as indicated by pink bars. As soon as the controller is turned off, the unpredicted errors caused by stepping itself accumulate and cause the pitch orientation to drift. Turning the controller back on restores the perceptual signal so that it matches the reference signal. Snapshot images to the right show trunk posture when the pitch controller is on and off. **(D)** Plots and snapshots have the same format as in (C), except that the roll controller is periodically turned off instead of the pitch controller. Roll control is also shown to effectively defend against the disturbances cause by stepping itself.

## Discussion

Here we present a legged robot capable of posture control and locomotion. The design is simple, consisting of a hierarchical network of negative feedback control system modules in which higher systems accomplish their goals by continuously adjusting the goals of lower systems. The design is also robust: the robot can reliably hit a sequence of position targets across rough and unstable terrain while successfully defending all of its goal states in parallel. The robot is able to do this with no prior knowledge of the environment or training, despite significant limitations in motor speed and power, transfer delays, and imprecise components.

The computational cost of this approach is very low. Our hierarchical control design does not require any explicit computation of inverse or forward kinematics and dynamics. Instead of performing inverse computations based on knowledge of detailed physical laws, and predicting how a pre-specified motor command will move effectors, the control modules in our are given top-down specification on *what to sense*. Their outputs will vary systematically with disturbances, defined as deviation from reference signals, to achieve their goals. This process is computationally simple, essentially involving calculating errors relative to goals and routing those errors to the appropriate lower systems. Moreover, since each degree of freedom is handled by a dedicated controller, increasing degrees of freedom only requires an increase in the size of the network without exponentially increasing the computational demand. Instead of forward computations, the output of every controller simply mirrors and opposes environmental disturbances relative to an internal goal value through subtraction. The essential computations can be implemented entirely by analog circuits, or by simple neural circuits of excitatory and inhibitory neurons.

### Comparison with other approaches

Because there is no general consensus on the operating principles of the nervous system, a wide variety of techniques has been used in neurally inspired robotics. One common approach is for a catalogue of predefined states or actions to be released by sensory or internal conditions. For example, a robot might exhibit stabilizing posture output in response to body tilt above a threshold value (4, 30, 46), or a modification of stepping output in response to a particular pattern of sensory input (28, 29, 36, 37, 41). A closely related approach is to convert sensory inputs into motor outputs in a continuous and graded manner using sensorimotor transformations (28). Other designs are entirely open loop and rely on internal network dynamics to generate behavior (35).

What all these designs neglect is how behavior is actively directed towards accomplishing a set of internally specified goals which are continuously changing; it is not simply ‘released’ or modified by environmental conditions. The goals or ‘set points’ can be adjusted by top down signals from higher levels (38). There is an explicit internal comparison between a sensory input and a goal value rather than a direct transformation of sensory input into motor output (52). Without this comparison and the resulting error, course correction towards a goal is not possible, and the value of lower reference states cannot be adjusted in service of hierarchically higher goals.

Although error reduction is widely used, for example in deep neural network training (31) and reinforcement learning (RL), error reduction in our robot is not used for learning but is instead responsible for generating behavior in real time. There are therefore fundamental differences between our approach and conventional approaches. For example, in a legged robot using RL, a readout map was trained through reward delivery to transform sensory inputs into modulatory changes to a hierarchically lower dynamic oscillator, such that a simulated robot could climb over difficult terrain (1). In other robots, RL is performed in simulation and the policy is then uploaded to a physical body to generate locomotion (13, 24, 32, 43). RL for locomotion has also been successfully performed online with no simulation (18). While the presence of a reward function gives these robots the appearance of goal-directedness, there is in fact no active pursuit of a behavioral goal, but instead only the shaping of a sensorimotor transformation to maximize reward. It may be argued that the reward function provides feedback based on a comparison between actual and desired robot behavior. However, such feedback is derived from the perspective of the observer or designer, and not related to the internal comparison as in our design. There is no internal definition of reward in RL. The externalist view of reward function is essentially open loop, as the loop can only be closed with the designer included, and the whole control architecture is not truly autonomous because it lacks any internal reference states. Consequently, there is no comparison of current input to the internal reference value, and thus no active course correction in approaching the goal. RL approaches usually require extensive offline training to establish the correct mapping of sensory input onto action output (19), whereas the present approach does not require any training because there is no state-action mapping.

A popular control architecture for legged robots, influenced by influential models of motor control in neuroscience (48, 49), is model-based control. This approach requires an accurate internal model of the robotic system. The state of this model is updated by sensory inputs and used to simulate its dynamics for the purpose of calculating the appropriate feedforward movement command. This command can be optimized according to a cost function defining the numerous goals of a task at a single level, such as moving the trunk of the robot to a desired position while also respecting contact configuration and friction constraints (13). The success of this approach is largely determined by the accuracy of the internal model (24). In robots with simple and well-characterized bodies, and environments with little or no unpredictable disturbances, model-based control can be successful. However, designing model-based control algorithms for legged locomotion is exceptionally challenging and requires sophisticated mathematics that cannot be computed by the nervous system (as opposed to scientists trained in symbol manipulation, familiar with physical laws, and using powerful computers capable of performing matrix calculations in real time). With increased degrees of freedom, model-based approaches present insurmountable computational challenges. In contrast, increasing degrees of freedom does not present significant challenges for our approach, as more controllers can simply be added at each level. No significant change is required, either in the basic algorithm used or in the design of the hierarchy.

Unlike conventional techniques outlined above, the control architecture presented here requires no learned policy or internal model of the body--only a representation of the relevant perceptual and goal values. Every module, with the exception of the burst generators, independently and continuously seeks to match its perception with its goal value through corrective outputs in real time. Unlike designs that only implement feedback control at the level of joint control and some form of feed-forward computation above that to generate behavior (11,14, 22, 25, 28, 32, 43, 45, 50), our design uses closed loop negative feedback control at every level of the hierarchy. This feature replicates the purposive nature of animal behavior, which is often mistakenly assumed to be a feedforward process whereby a stimulus input is transformed by the nervous system and results in motor output (51).

Each controller simply calculates the errors between actual and desired perceptions and convert the errors into outputs that reduce the same errors. The major computation involved is performed by a comparison function that subtracts one signal from another. Higher goals are achieved by specifying the goals to be achieved by lower levels. This design is characterized by parallel and decentralized processing, as well as from the ability of a hierarchy to collapse and simplify complex tasks into a hierarchy of simpler subgoals.

Although the benefits of hierarchical architectures have long been known, there is no known framework for how they might be implemented biologically. For example, a recent review acknowledges the many advantages of hierarchical motor control, but admits there is no plausible model for modular control and communication between modules and levels in the nervous system (34). Here we propose such a model.

Our design is directly inspired by the hierarchical organization of the motor system in vertebrates (16), and by its ability to pursue a hierarchy of goals in parallel, despite a complex and unpredictable environment (52). This basic design is in agreement with extensive experimental work in neuroscience (5, 6, 7, 8, 20, 21, 26, 27), and the efficacy of this design supports input control hierarchies as a working model for how the nervous system generates goal-directed behavior.

Our design is only a prototype with only 4 levels in the hierarchy, but it is not difficult to add higher levels and additional sensors to generate higher forms of goal directed behavior. Our design can also permit unsupervised learning, defined as changes in system parameters. Such learning can take place in a distributed fashion, with no need for backpropagation techniques often used in conventional neural networks.

In summary, the network architecture presented here, inspired by the vertebrate nervous system and tested with a working robot, suggests a new and powerful approach in robotics. It provides a foundation for future work building testable models of the nervous system as well as in the development of intelligent systems more generally.

## Materials and methods

### Apparatus

The robot is a legged quadruped with 3 motor actuated joints per leg. The body is made from 3D printed parts and light aluminum channeling. Motors are standard size hobby servos (HS-7954SH, Hitec) with gear drives to increase strength (Servo City). Knees used a 2:1 gear reduction while shoulder motors used a 3.8:1 gear reduction. These joints are not easily back-drivable. The body weighs 7.8 kg and measures approximately 60cm long by 40cm wide. Upper leg joints are located 45 cm apart lengthwise and 19 cm apart widthwise. To help support the body weight and reduce gear play, a torsion spring is installed in each joint to oppose the effect of gravity. Torsion springs transmit force by gears into the drive train of each joint. Torsion gears add force in parallel with servomotors.

Each joint is equipped with an analog position sensor and an analog torque sensor between the motor and the movable body part. Joint angular velocity is sensed by taking the first derivative of joint position from one step of the control loop to the next. Each foot is equipped with an analog force sensor for detecting ground reaction forces. For a vestibular system, the trunk of the robot houses an analog Inertial Measurement Unit (IMU) consisting of a 2-axis accelerometer and two gyros for sensing orientation of the body in space. Sensory inputs are communicated from the body to the computer by a shielded multi conductor cable and read into a standard desktop PC as analog voltages through a National Instruments Box (NI USB-6225). Motor outputs are commanded from the PC through a USB servo control chip (Maestro 24, Pololu). There is a latency of approximately 75 ms between a command being issued and movement by the motor. The robot uses a tethered power supply (7.4 V and 20a).

Torque is controlled by converting the torque controller output into a motor position through a virtual driven damped torsion spring, a feature which was included to smooth outputs and improve compliance to small high frequency disturbances. Although virtual spring compliance makes the behavior of effectors more dynamic and unpredictable, this does not pose a problem for our approach which reduces perceptual errors regardless of their source. Damping constant and spring constant values were selected to generate a stiff spring system exhibiting small compressions relative to the overall movement of the joint. The spring has a resting length of zero. The virtual spring is represented by the following equation:

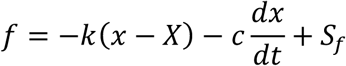

Here, *X* is the position of the virtual driver determined by the output of the torque controller. *x* is the position of the virtual mass connected by a spring to the driver, which determines the servomotor reference value. By adjusting *X*, we adjust the length of the virtual spring and thus the force acting on *x*.. *k* is the spring constant, *c* is the damping constant, and *S*. is the force sensed by the torque sensor. The total force, *f*, acting on the effector at a given time determines the acceleration of *x*.

Cartesian coordinates are calculated from joint angles and segment lengths using trigonometry. For each leg, we calculate the foot and knee position in Cartesian space with the shoulder as the origin. To position these body parts in egocentric space, we then translate these values by adding or subtracting fixed offsets along the horizontal plane so that each shoulder is positioned the correct measured distance from the origin at the center of the trunk. To achieve position relative to gravity, we then rotate all body points around the origin according to the pitch and roll angles sensed by the IMU.

The controllers operate according to the following equations:

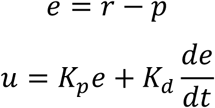

Here, *e* is error, *r* is reference signal, and *p* is perceptual signal, *o* is output signal, *K*_*p*_ is proportional gain, and *K*_*d*_ is derivative gain.

The gain scheduler interfacing between level 3 (posture) and level 2 (joint angular velocity) systems functions to dynamically scale the outputs from level 3 systems to produce posture velocities that retain their linear relationship with those outputs regardless of the current body posture. This feature is not essential; as long as the PD gains are appropriate, and as long as we convert a given posture error into a set of joint velocity reference signals which oppose that error, then the error will approach zero (Powers, 2005). However, gain scheduling is useful for stabilizing large and fast outputs during locomotion. The gain scheduler works as follows: from the current perceived body posture, PD outputs from active level 3 controllers are converted into movements in the opposite direction as the errors that caused them, to generate a new ‘virtual’ posture. This new posture is then subtracted from the current perceived posture to calculate the joint angle changes between them. These angles determine proportionate velocity reference signals sent to corresponding level 2 joint angular velocity controllers.

Once the network was assembled, controller PD gains were tuned to accomplish a good balance of stability and speed. When tuning a layer of the hierarchy, the higher layers were left inactive and the reference signals were adjusted by the experimenter to assess performance. The key performance standard for the network as a whole is the ability to reliably reach locomotor targets through an environment containing unknown disturbances. Secondary performance standards include the successful defense and pursuit of lower reference states during locomotion, including trunk and foot positioning and joint control.

### Testing controllers

To test foot position controllers, the robot was suspended hanging and tipped by the experimenter while foot controllers for a single leg were active. To test trunk orientation controllers, the robot was placed, with all trunk position and orientation controllers active, on a platform which was randomly tipped by the experimenter along pitch and roll planes. To test the viability of the proposed network architecture for locomotion, the robot was required to navigate across terrains of unstable and loose and piled rocks to achieve a sequence of targets. Piled rock obstacles formed sloped terrains with slopes of between 5 and 20 degrees. The robot was able to complete successful crossings over diverse terrains while successfully defending all controlled variables.

## Acknowledgements

We would like to thank Konstantin Bakhurin and Ryan Hughes for helpful discussion during the writing of this paper and comments on the manuscript.

## Movies

**Movie 1. Disturbance to trunk pitch and roll**. The robot is standing on a platform which is randomly tilted by the experimenter. Pitch and roll reference signals are set to zero and do not change.

**Movie 2. Voluntary trunk pitch and roll movement**. The robot is standing while the trunk pitch and roll reference signals are randomly adjusted by the experimenter. Internal plots of perceptual and reference signals for those systems is also shown.

**Movie 3. Voluntary trunk up-down and yaw movement**. During this Movie, the robot is standing while the experimenter randomly moves a joystick to change trunk and yaw reference signals.

**Movie 4. Yaw orientation during stepping**. The yaw reference signal is adjusted while the robot is stepping. As the legs continuously reset, trunk yaw reference signal change during stepping results in controlled whole-body orientation change.

**Movie 5. Locomotion in a straight line across flat terrain**. The trunk X reference signal is adjusted during stepping, resulting in controlled translation of the whole body in space across flat terrain.

**Movie 6. Resisting disturbances during stepping**. The robot successfully steps in place without falling while push disturbances are introduced.

**Movies 7. Sequenced target position control through yaw orientation and locomotion across flat terrain**. The robot is given an arbitrary sequence of three position targets which it pursues one at a time through locomotion. For a given sequence step, orientation towards the target is controlled by adjusting the yaw reference signal during stepping, while distance to target is controlled by adjusting the egocentric X position reference. A wireframe recording of the robot’s internal perceptions are also shown. The red dot indicates the target position.

**Movie 8. Sequenced target position control through yaw orientation and locomotion across loose rocks**. Sequence control task is identical to that shown in Movie 7. Two sessions are shown with different terrains including loose rocks.

**Movie 9. Sequenced target position control through yaw orientation and locomotion across piled rocks**. Sequence control task is identical to that shown in Movies 7-8. Two sessions are shown with different terrains including piled and loose rocks.

